# Forecasted trends of the new COVID-19 epidemic due to the Omicron variant in Thailand, 2022

**DOI:** 10.1101/2022.01.24.477479

**Authors:** Rapeepong Suphanchaimat, Pard Teekasap, Natthaprang Nittayasoot, Mathudara Phaiyarom, Nisachol Cetthakrikul

## Abstract

**Background:** The introduction of the Omicron variant is of significant concern to the Thai Government due to the possibility of a new wave of the COVID-19 epidemic, which may cause a huge strain to the country’s health system. This study aims to forecast the trends of COVID-19 cases and deaths given the advent of the Omicron variant in Thailand.

**Methods:** We used a compartmental susceptible-exposed-infectious-recovered model in combination with a system dynamics model. We developed four scenarios according to differing values of the production number (R) and varying vaccination rates.

**Results:** The findings indicated that in the most pessimistic scenario (R = 7.5 and base vaccination rate), the number of incident cases reached a peak of 49,523 (95% CI: 20,599 to 99,362) by day 73 and the peak daily deaths enlarged to 270 by day 50 (95% CI: 124 to 520). The predicted cumulative cases and deaths at the end of the wave (day 120) were approximately 3.7 million and 22,000 respectively. In the most optimistic assumption (with R = 4.5 and a speedy vaccination rate [tripled the base rate]), the peak of the incident cases was about one third of the most pessimistic assumption (15,650, 95% CI: 12,688 to 17,603). The corresponding daily fatalities were 72 (95% CI: 54 to 84) and the prevalent intubated cases numbered 572 (95% CI: 429 to 675).

**Conclusions:** In the coming months, Thailand may face a new wave of the COVID-19 epidemic due to the Omicron variant. The case toll due to the Omicron wave is likely to outnumber the earlier Delta wave, but the death toll is proportionately lower. Despite the immune-escape characteristic of the Omicron variant, the vaccination campaign for the booster dose should be expedited as an effective way of preventing severe illness and death.

## Introduction

Over the past few years, the world has recognized the Coronavirus disease 2019 (COVID-19) pandemic as one of the most serious health threats in human history. The disease is caused by Severe Acute Respiratory Syndrome Coronavirus-2 (SARS-CoV-2), which is transmitted by direct contact with droplets containing pathogens and indirect contact with contaminated surfaces [1, 2]. The first reported case of COVID-19 was found in China, then the disease spread rapidly throughout the world and became a global pandemic [3]. At the time of writing, the global case toll has reached almost 300 million with approximately 5.5 million accumulated deaths [4].

A SARS-CoV-2 variant with a significant genetic change from the original strain which is demonstrated to have the following changes at a degree of global public health significance (namely, increase in transmissibility, enhancement of clinical virulence, and decrease in effectiveness of public health measures or vaccines) is characterized by the World Health Organization (WHO) as a variant of concern (VOC) [5, 6]. So far, the VOC comprises Alpha variant (B.1.1.7), Beta variant (B.1.351), Gamma variant (P.1), Delta variant (B.1.617.2), and Omicron (B.1.1.529) [6]. During the second and the third quarters of 2021, the world was severely hit by the Delta variant which was firstly detected in India. In June 2021, the WHO indicated that the Delta variant had become the dominant strain globally. It is believed to be responsible for the deadly second wave in India, and, subsequently, it drove a steep rise of daily infections in many parts of Asia and the US [7, 8].

By late 2021, while the world was hoping to see a promising end to the pandemic as global incident cases gradually subsided, another global threat, the Omicron variant, was reported. It was believed to have numerous mutations with the potential to increase transmissibility compared with prior variants, and to partially escape infection- or vaccine-induced immunity [9–11]. The variant was responsible for numerous clusters in South Africa, and it now spreading widely to more than 110 countries (as of 1 January 2022) [12]. Many countries in Europe and the UK are now witnessing a day-to-day new record high of COVID-19 cases due to the Omicron variant.

Thailand is among many countries that have been critically affected by the COVID-19 pandemic. The first COVID-19 wave in Thailand occurred during March-May 2020 due to super-spreading events from a boxing stadium and a nightlife hotspot in Bangkok downtown [13]. The second wave originated from a cluster of cases in a shrimp market in the inner city of Samut Sakhon and lasted between December 2020 and February 2021 [13, 14]. The third wave was mostly caused by the Alpha variant in April 2021, followed by the fourth wave beginning in June 2021 due to extensive local transmission of the Delta variant [14]. During that time, the Thai Government implemented a lockdown policy to mitigate the magnitude of cases and deaths as a pre-emptive measure to avoid the collapse of the healthcare system. Almost all international flights were banned and all inbound travellers were obliged to undergo a 14-day quarantine upon arrival. COVID-19 vaccines, both by importation and domestic production, were rapidly rolled out. By early December, the volume of Thais receiving at least one shot of the COVID-19 vaccine numbered about 70% of the total population; the benchmark believed to make the country achieve herd immunity as per the national vaccine plan of the Government [15]. By the end of 2021, Thailand saw a case toll of 2.2 million and approximately 22,000 fatalities.

The number of new daily cases and deaths in Thailand reached its peak by mid-August 2021 (about 23,000 cases and 290 deaths a day). Then, the incident case volume continuously subsided to a level below 3,000 by mid-December 2021 [16]. As the situation appeared to be relieved, the Government later withdrew the lockdown policy by November 2021 but still encouraged people to keep physical distancing and maintaining mask wearing in public spaces. To prepare for the resuming of international flights to boost the country’s touristy businesses, the Government planned to implement a “Test & Go” policy in which an inbound traveler is not required to undertake a 14-day stay in the quarantine center as long as he/she is fully vaccinated and possesses a proof of evidence showing negative SARS-CoV-2 detection (by reverse transcription polymerase chain reaction [RT-PCR]) 72 hours prior to departure and again presents with a negative result by RT-PCR upon arrival [17].

However, by early December 2021, the Thai Ministry of Public Health (MOPH) declared the discovery of the first imported case of the Omicron variant. A few weeks later, the Government pre-emptively suspended the Test & Go policy in order to block the potential importation of Omicron cases despite the fact that by mid-December 2021, the local transmission of the Omicron variant was confirmed. The situation caused concern for the Government because the Omicron variant could create a serious threat to the Thai healthcare system similar to that during the Delta pandemic. This point informs the objective of this study.

This study aims to forecast the trends of new cases as well as the death toll and use of health resources for severe cases given the advent of the Omicron variant in Thailand. We hope that the findings of this study will help aid policy decisions for optimal preparation of the healthcare system resources, and highlight the importance of measures (including vaccines and non-pharmaceutical interventions [NPI]) which may help mitigate the outbreak magnitude.

## Methods

### • Study design

A secondary data analysis was employed. Most parameters used in this study were acquired from the internal database of the Department of Disease Control (DDC) and the Department of Medical Services (DMS), the MOPH. Some basic parameters, such as incubation period and infectious duration of infection were obtained from international literature. Parameters reflecting the Thai healthcare system performance were obtained from expert opinions and model adjustment. The forecasting duration was 120 days. It is worth-noting that as the Omicron variant is quite new to the world, some variant-specific parameters are not available at the time of writing. We therefore adopted the parameters specific to the Delta variant instead.

### • Model framework

We employed a compartmental susceptible-exposed-infectious-recovered (SEIR) model and the system dynamics (SD) model to frame the analysis [18, 19]. The simplified model framework is demonstrated in Fig 1. We divided the entire Thai population into four groups based on the vaccination profile: (i) the unvaccinated, (ii) the one-dose, (iii) the two-dose, and (iii) the booster (receiving at least three shots of vaccine). In each group, we sub-categorized the population into five sub-categories according to the infection status: (i) the susceptible, (ii) the exposed, (iii) the infectious before isolation, (iv) the infectious after isolation, and (v) the recovered.

**Fig 1.**
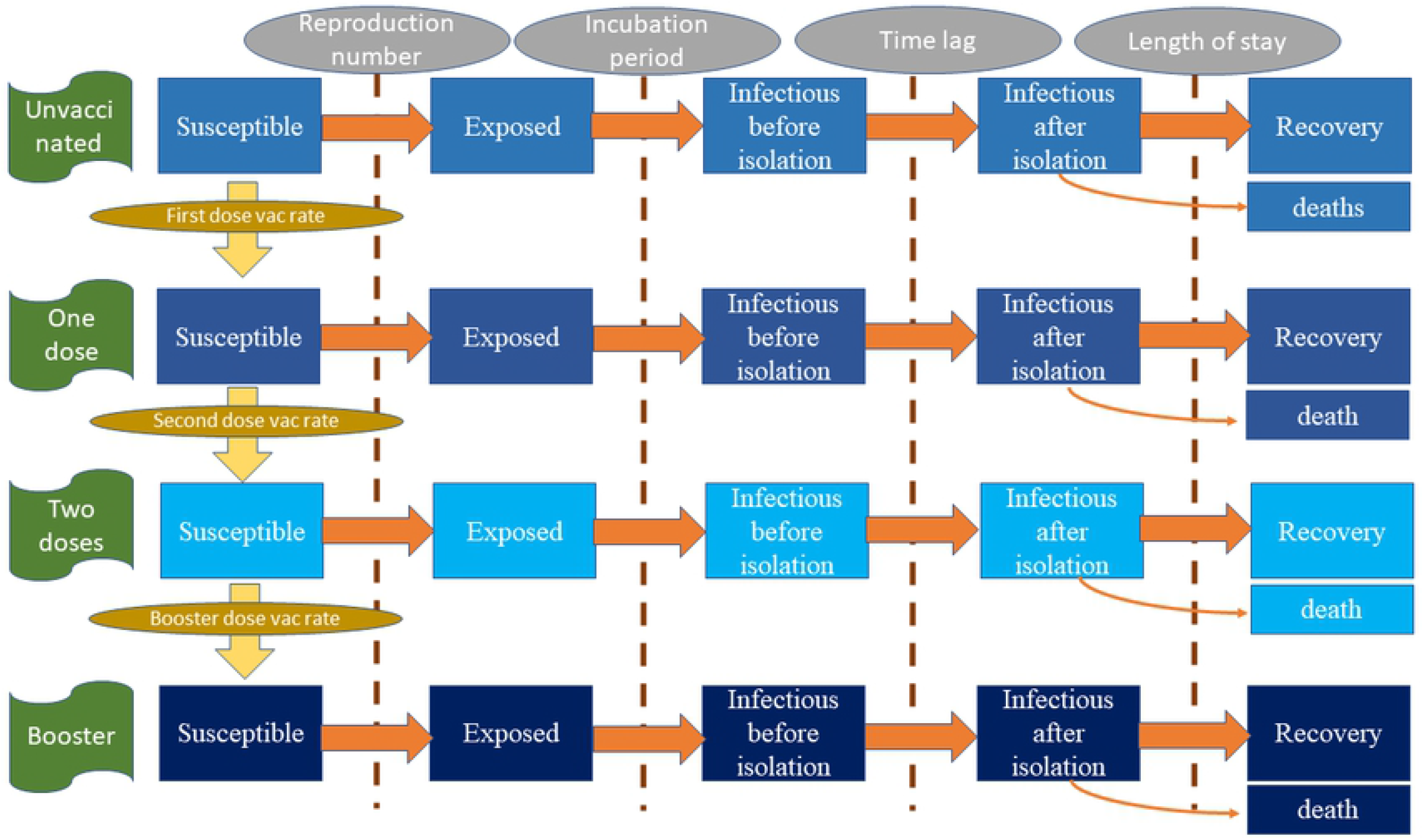
Model framework.

The speed of transfer from susceptible group to exposed group was mainly influenced by the reproduction number (R) [20]. The transition from the exposed group to the infectious group depended on the incubation period. We adapted the traditional SEIR model by splitting the infectious group into before isolation and after isolation. The reason behind this is that once admitted to a hospital, an infected person would be isolated by the hospital protocol (suppose no nosocomial infection). The length of stay (LOS) in a hospital influenced the speed of recovery. Among the admitted patients, the prevalence of intubated cases attracted the attention of policy makers the most. This is because the volume of intubated cases represent the reserve capacity of intensive care, while asymptomatic or mild cases are allowed to be isolated at home or in the community according to the current MOPH protocol [21]. We further assumed that some of the intubated cases later died and no deaths occurred without intubation. An unvaccinated susceptible person encountered two paths, either becoming exposed to the disease or remaining as a susceptible person and receiving the first vaccine shot, which depended on the vaccination rate in the entire population. The same concept also applied for the one-dose, the two-dose, and the booster groups.

### • Model assumptions, parameters and formula

The model was governed by the following assumptions. First, the population was homogenously mixed, inferring that all susceptible individuals were subject to infection (with varying probabilities conditional on vaccination status). This assumption coincided with present evidence which pointed to the immune-escape property of the Omicron variant [22, 23].

Second, in most pandemics, the exact number of initial infectees could be hardly identified. We proposed that the volume of infectees equaled 10,000; about threefold the size of daily incident cases at the time of writing. This assumption corresponded with the experience of the outbreak investigators of the DDC that when a super-spreading event was notified, approximately three to four generations of infection had already passed by.

Third, there existed some degree of under-reporting for asymptomatic and mildly symptomatic infectees. Literature suggested that the underreporting phenomenon was commonly found in numerous countries during the surge of the outbreak [24]. The field operation of the Rural Doctor Society of Thailand in mid-2020 affirmed that, from an active screening, over 13% of people in high-density communities in Bangkok were found infected with COVID-19 but their records were not present in the official infectee list of the MOPH [19]. However, we postulated that there was no underreporting of intubated cases and deaths.

Fourth, recent evidence affirmed that the Omicron variant transmitted more easily and faster than the Delta variant. Based on the model calibration against the incident cases during the peak of the Delta epidemic (July-December 2021), we found the value of 1.38 best represented the R for the entire population. We further proposed that if the Omicron variant caused another epidemic wave in Thailand, its R value would be about 3.1-5.4 times larger than that of the Delta variant [25, 26]. These figures were later used to construct the model scenarios.

Fifth, in reality, the R did not remain constant over time due to social adaptability and various NPI. We postulated that it took 15 days for the R of Omicron variant to climb from the current R in Thailand at the time of writing (0.86) to reach the set value (fourth assumption), then it naturally dropped by two points within 60 days later.

Fifth, the R was influenced by two key factors: (i) the vaccine effectiveness (VE) against any infection, and (ii) the contact rate of the people in the society. The COVID-19 vaccine acted not only on the probability from being susceptible to being exposed but also altered the severity profile of the infectious compartment (reducing the probability of becoming severe cases or deaths). Since no officially published report of the VE against the Omicron infection in Thailand had come out yet, we used the VE of the viral-vector vaccines in the UK instead (also the same vaccine type widely administered in Thailand) [27].

Sixth, we used the vaccination rate by late 2021 as a base vaccination rate in the population and assumed that this remained unchanged throughout the study course. We touched upon the vaccination rate again in the later section, “Model scenarios and interested outcomes”. The COVID-19 vaccines were administered only when an individual lay in the susceptible state.

Last, linked to the fifth assumption, the contact rate of individuals depended on the magnitude of the outbreak. Social measures and individual protective behaviors become stricter when the volume of incident cases enlarges. These factors were also considered as part of the NPI. This idea concurred with the fact that the mobility trend (using public transport use as a proxy) of Thai individuals diminished by 68% during the Delta epidemic, compared with the pre-COVID-19 era. By December 2021, when the Delta epidemic declined, the mobility trend declined by 25%, relative to before 2020 [28]. We therefore added a parameter reflecting the NPI effectiveness in the model.

We used Microsoft Excel and Stella 2.0 (number: 251-401-786-859) for model execution. Tables 1–2 exhibit important parameters and formulas of the model.

**Table 1.** List of essential parameters.

| Parameters | Unit | Value | Reference (note) |
| --- | --- | --- | --- |
| Reproduction number | Unitless | 4.3-7.5 | Ito et al. [25], Head and van Elsland [26] (3.1- 5.4 times greater than the Delta epidemic in Thailand in 2020) |
| Population | Persons | 66.2 * 10<sup>6</sup> | National Statistical Office of Thailand [29] (assume homogenous mixing) |
| Mean infectious duration | Days | 4.6 | Hart et al. [30] (assume gamma distribution with scale parameter of 0.03 and shape parameter of 165.9—same as the Delta variant) |
| Mean incubation period | Days | 3.2 | Helmsdal et al. [31] (assume gamma distribution with scale parameter of 0.01 and shape parameter of 302.7—shorter than the Delta variant) |
| Time lag from being infected to isolation | Days | 5 | Model calibration (assume same as the Delta epidemic in Thailand in 2020) |
| Initial number of infectees | Days | 10,000 | Model calibration (assume fourfold greater than the present incident cases) |
| Initial proportion of unvaccinated population | Unitless | 0.26 | Internal database of the Department of Disease Control |
| Initial proportion of one-dose vaccinees | Unitless | 0.09 | Internal database of the Department of Disease Control |
| Initial proportion of two-dose vaccinees | Unitless | 0.58 | Internal database of the Department of Disease Control |
| Initial proportion booster-dose vaccinees | Unitless | 0.06 | Internal database of the Department of Disease Control |
| First-dose vaccination base rate | Persons/day | 66,200 | Internal database of the Department of Disease Control (assume equaling the rate of booster dose) |
| Second-dose vaccination base rate | Persons/day | 198,600 | Internal database of the Department of Disease Control (the largest rate compared to other doses as the second shot currently being the main policy priority) |
| Booster-dose vaccination base rate | Persons/day | 66,200 | Internal database of the Department of Disease Control |
| Vaccine effectiveness against any infection for one-dose vaccination | Unitless | 0.17 | Adapted from Head and van Elsland [26] and internal database of the Department of Disease Control |
| Vaccine effectiveness against any infection for two-dose vaccination | Unitless | 0.41 | Adapted from Head and van Elsland [26] and internal database of the Department of Disease Control |
| Vaccine effectiveness against any infection for booster-dose vaccination | Unitless | 0.65 | Adapted from Head and van Elsland [26] and internal database of the Department of Disease Control |
| Vaccine effectiveness against severe infection for one-dose vaccination | Unitless | 0.70 | Adapted from Head and van Elsland [26] and internal database of the Department of Disease Control |
| Vaccine effectiveness against severe infection for two-dose vaccination | Unitless | 0.90 | Adapted from Head and van Elsland [26] and internal database of the Department of Disease Control |
| Vaccine effectiveness against severe infection for booster-dose vaccination | Unitless | 0.95 | Adapted from Head and van Elsland [26] and internal database of the Department of Disease Control |
| Proportion of intubated cases to existing active infectees | Unitless | 0.005 | Internal database of the Department of Disease Control (assume half of the proportion of the Delta variant) |
| Ratio of deaths per existing intubated cases | Unitless | 0.03 | Internal database of the Department of Disease Control (assume same as the ratio of the Delta variant) |
| Length of stay for non-intubated cases | Days | 10 | Internal database of the Department of Disease Control |
| Length of stay for intubated cases | Days | 21 | Internal database of the Department of Disease Control |
| Non-pharmaceutical intervention base effectiveness against any infection | Unitless | See supporting information (S1 Fig) | Assume being an exponential function with the incident cases |

**Table 2.**
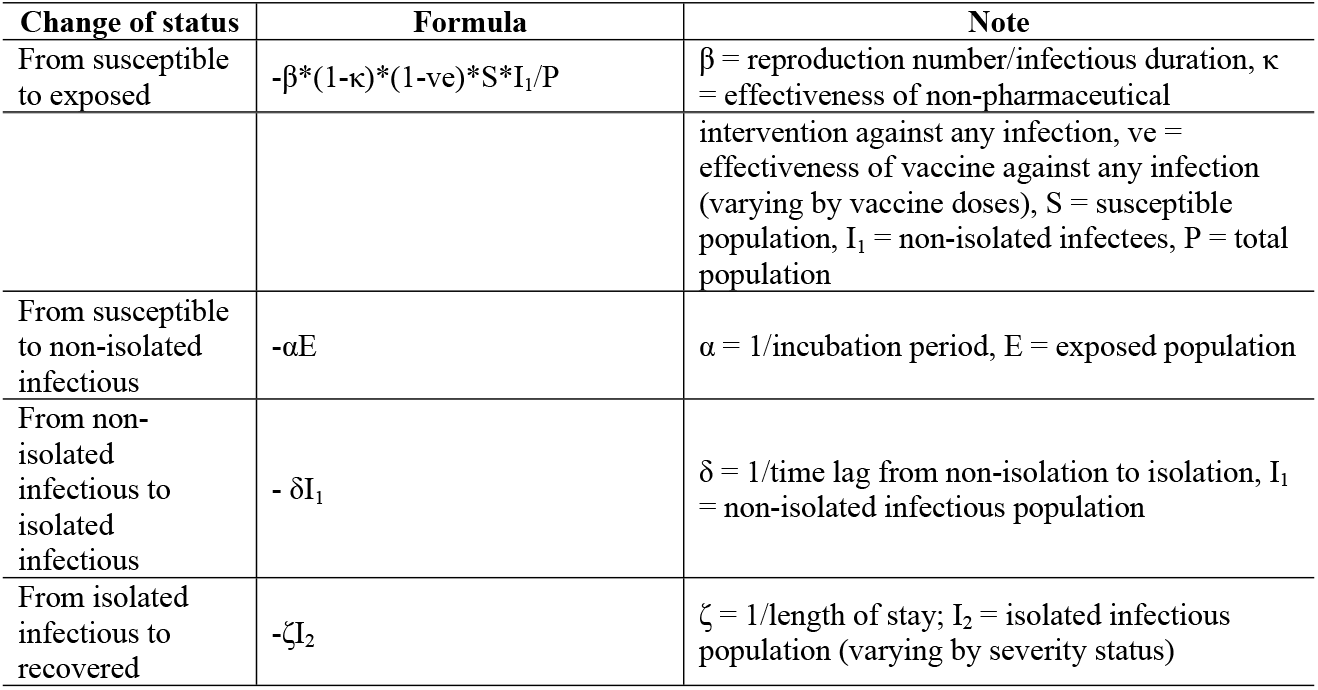
Essential formula of the model.

### • Model scenarios and interested outcomes

We focused on the following outcomes: (i) daily reported incident cases, (ii) daily deaths, (iii) prevalent intubated cases (requiring invasive ventilator), (iv) cumulative case toll and (v) cumulative death toll. These outcomes were commonly used by the MOPH to gauge the healthcare burden. We constructed the interested scenarios by varying the values of R (highly transmitted [R = 4.3] versus very highly transmitted [R = 7.5]) and the vaccination rate (base pace [as shown in Table 1] *versus* speedy pace [three times faster than the base pace]), Table 3. Scenario 1 was most pessimistic whereas scenario 4 was most optimistic. Scenarios 2-3 were between the two ends. The model was run for 100 times for each scenario, using infectious duration and incubation period as sensitivity parameters.

**Table 3.** Scenarios of interest.

| Scenario | Reproduction number | Vaccination rate |
| --- | --- | --- |
| 1 | 7.5 | Base pace |
| 2 | 7.5 | Speedy pace |
| 3 | 4.3 | Base pace |
| 4 | 4.3 | Speedy pace |

### • Ethics consideration

As this study used only secondary data available in the international literature and the database of the DDC and did not involve human participation, ethics approval was not required. Yet, we strictly conformed with the principles of ethical standards stipulated in the Declaration of Helsinki. The findings were presented in a way to prevent identifying an individual.

## Results

We first presented the number of estimated daily reported cases in Fig 2. In the most pessimistic scenario (scenario 1), the peak daily incident cases exceeded other scenarios. The incident cases would reach 49,523 per day, by day 73 (95% CI: 20,599 to 99,362). Scenario 2 where the vaccination rate was sped up but the R remained high (7.5) saw the peak incident cases of 30,025 (95% CI: 19,358 to 54,317). With the R dipping down to 4.3 in scenarios 3 and 4, we found the peak of the daily new cases at 16,889 (95% CI: 14,644 to 18437) and 15,650 (95% CI: 12,688 to 17,603) respectively by about day 50.

**Fig 2.**
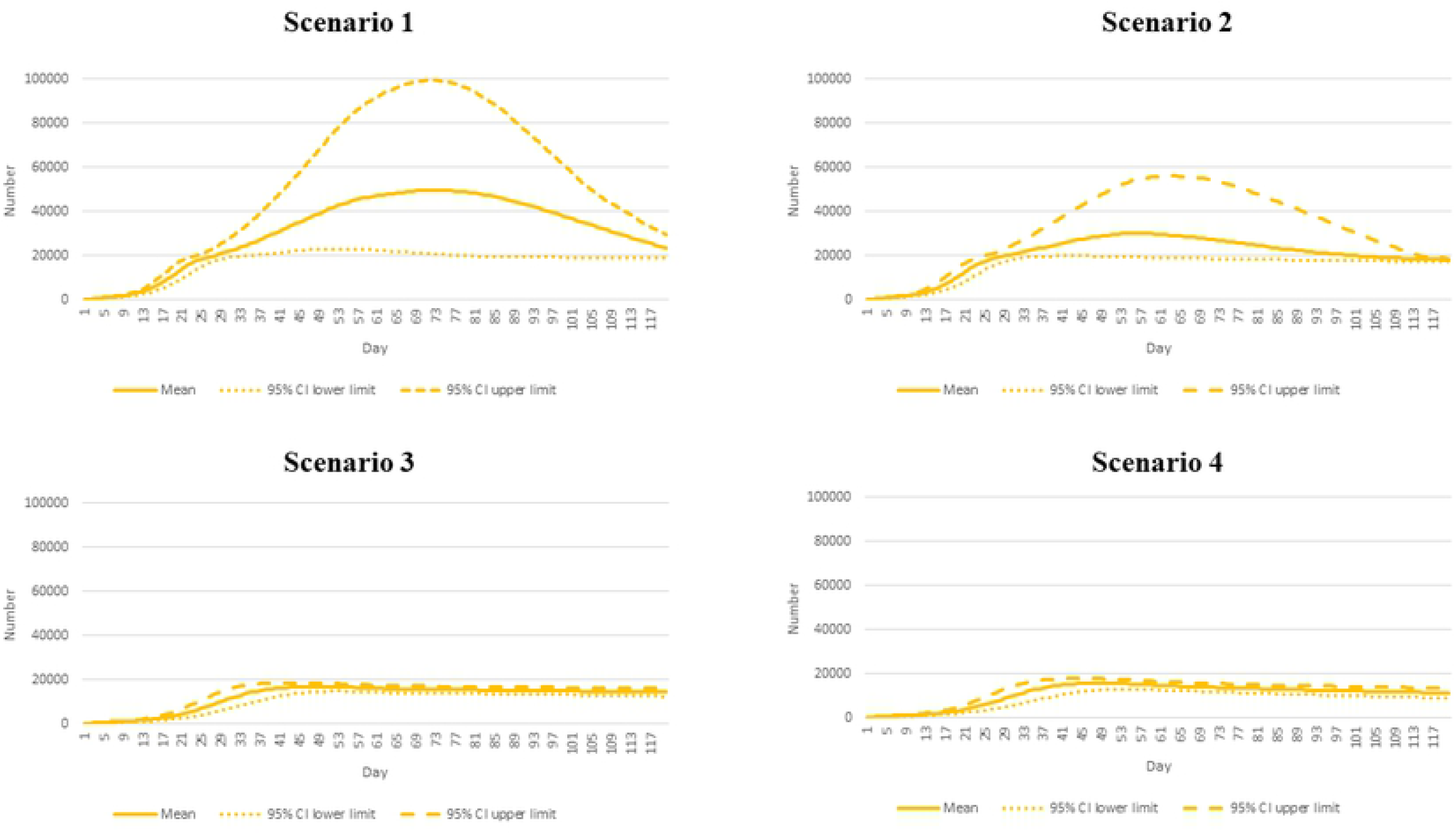
Daily incident cases by different epidemic scenarios.

For daily deaths, the number exceeded 100 by day 5 and reached the peak at 270 (95% CI: 126 to 518) by about day 50 in scenario 1. The second highest toll presented in scenario 2 with a peak of 129 deaths by approximately day 30 (95% CI: 91 to 211). Scenarios 3 and 4 demonstrated almost the same pattern over the course of the analysis. Scenario 4 showed the smallest number of peak daily deaths relative to other scenarios (72 deaths by day 28, 95% CI: 54 to 84), Fig 3.

**Fig 3.**
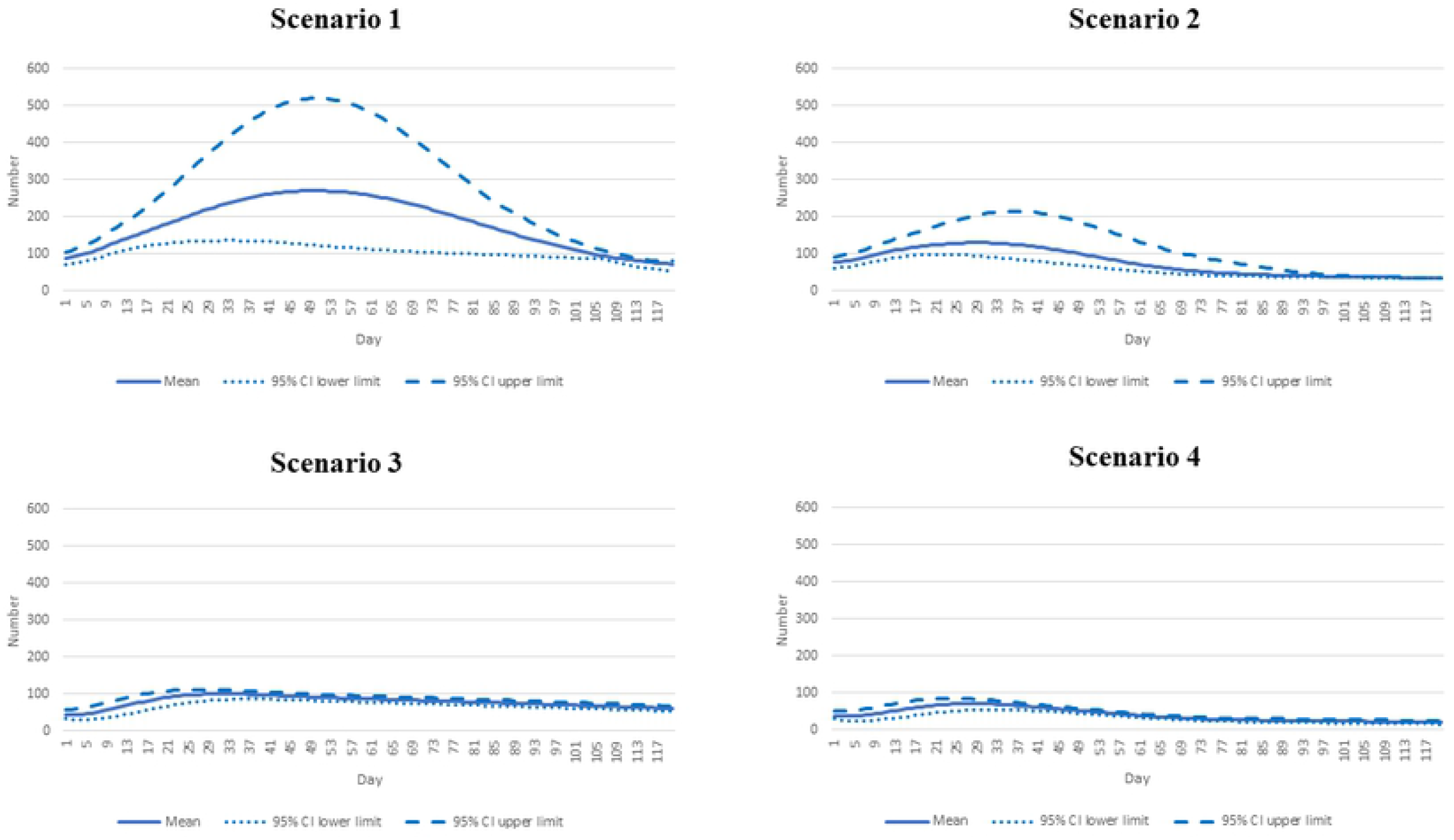
Daily deaths by different epidemic scenarios.

The prevalence of intubated cases followed the same pattern as daily deaths. Scenario 1 saw the highest number of prevalent intubated cases (2,161 cases by 50, 95% CI: 987 to 4,161). Scenario 2 exhibited the second highest peak, following scenario 1. The peak number reached 1,035 by day 30 (95% CI: 743 to 1,658). In scenario 3, the peak dropped to 800 during days 31-32 (95% CI: 675 to 888) and this further dipped to 572 in scenario 4 (95% CI interval: 429 to 675), Fig 4.

**Fig 4.**
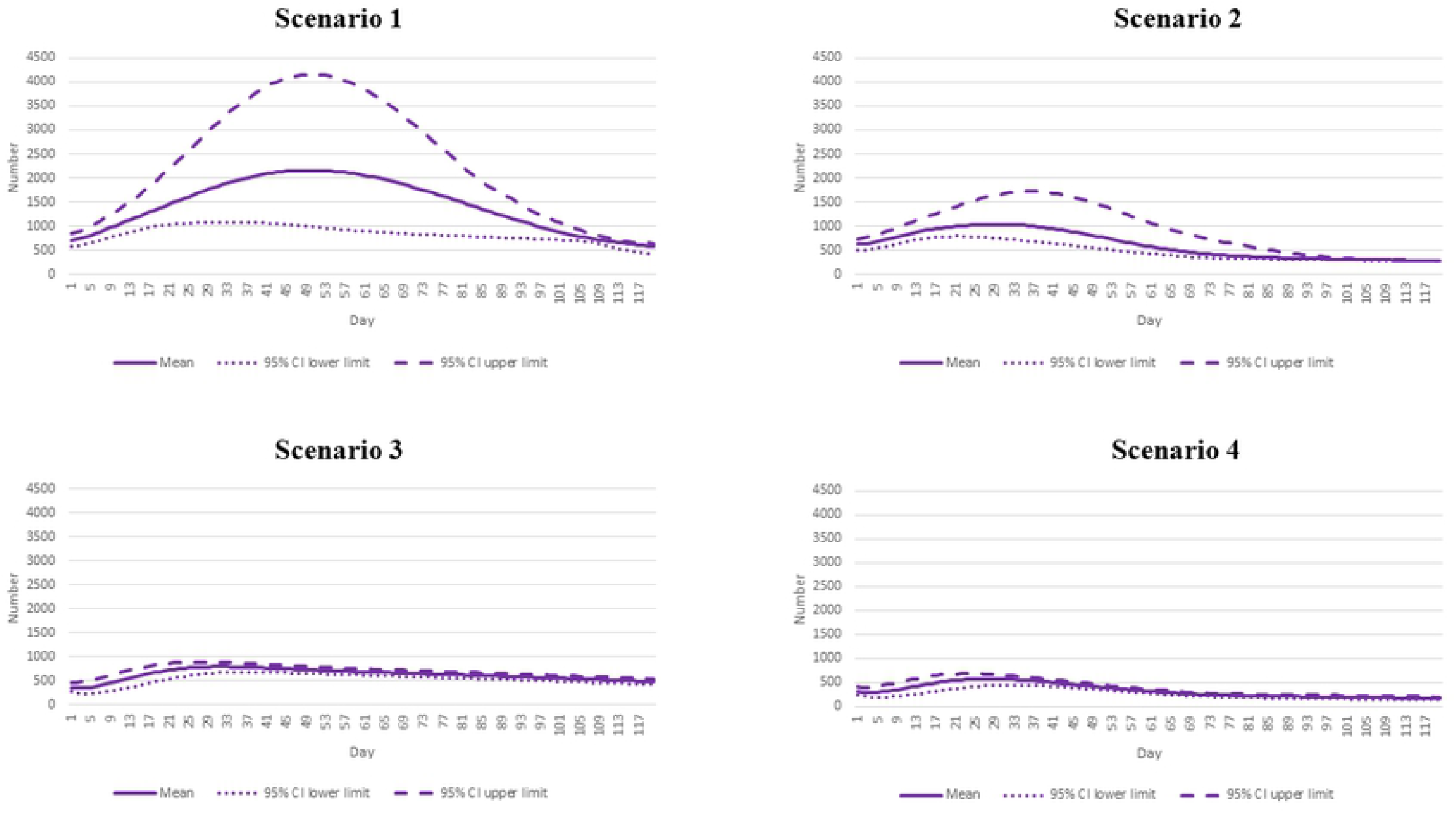
Prevalent intubated cases by different epidemic scenarios.

The mean of cumulative incident cases by day 120 accounted for approximately 3.7 million (95% CI: 2,035,674 to 6,401,118). This number was about threefold higher than the mean cumulative cases in scenario 4 (about 1.3 million, 95% CI: 1,016,258 to 1,521,434). The mean cumulative case toll in scenarios 2-3 was between 1.5 million and 2.4 million, Fig 5.

**Fig 5.**
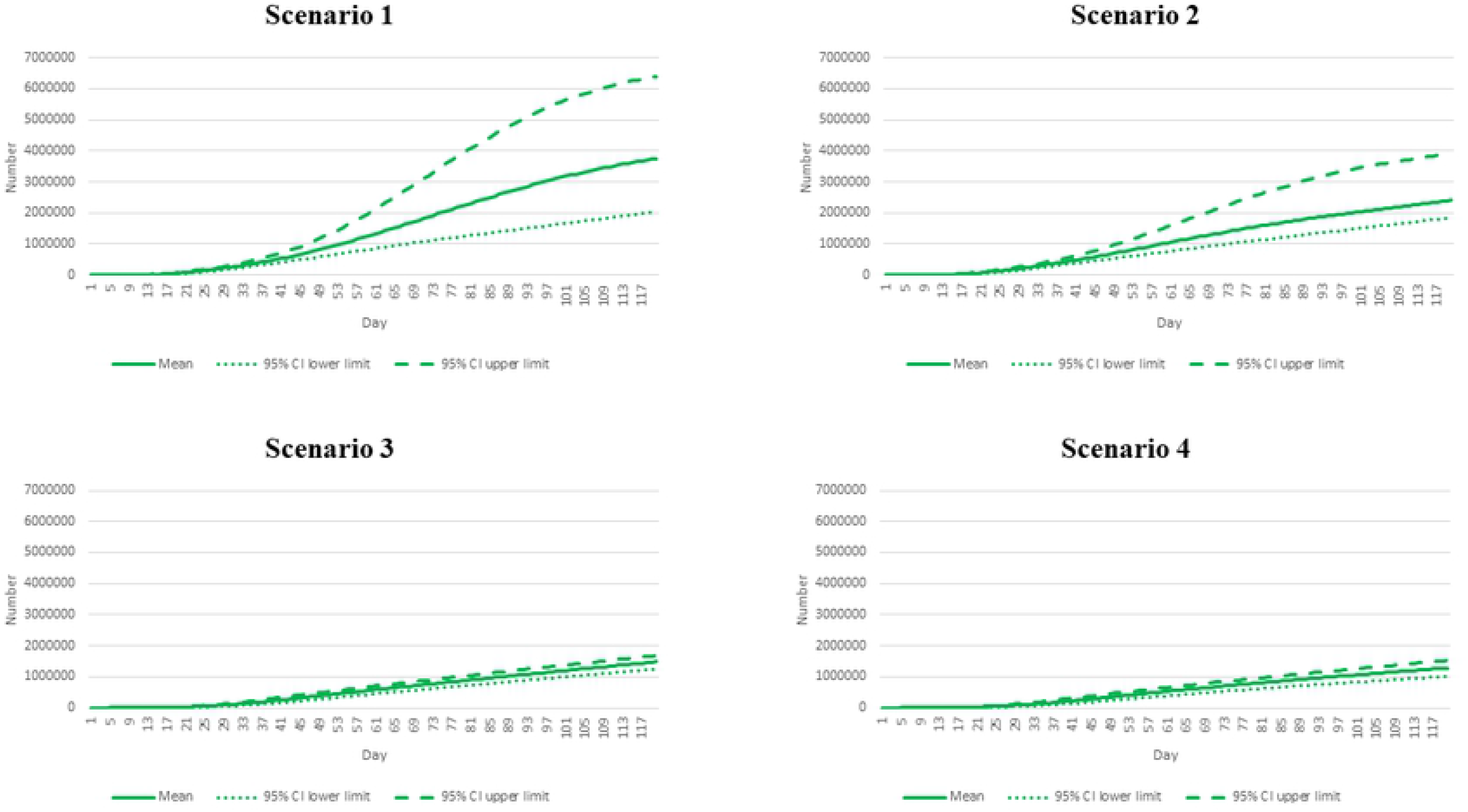
Cumulative case tolls by different epidemic scenarios.

Like other indicators, by day 120, scenario 1 presented the largest mean cumulative deaths at 21,649 (95% CI: 12,812 to 34,566). The mean death toll in scenario 4 was 4,996 (95% CI: 3,922-5,948), just a quarter of the size of scenario 1’s figure. Scenario 2-3 showed almost the same volume of cumulative deaths at between 9,163 and 9,467 by the end of the study period, Fig 6.

**Fig 6.**
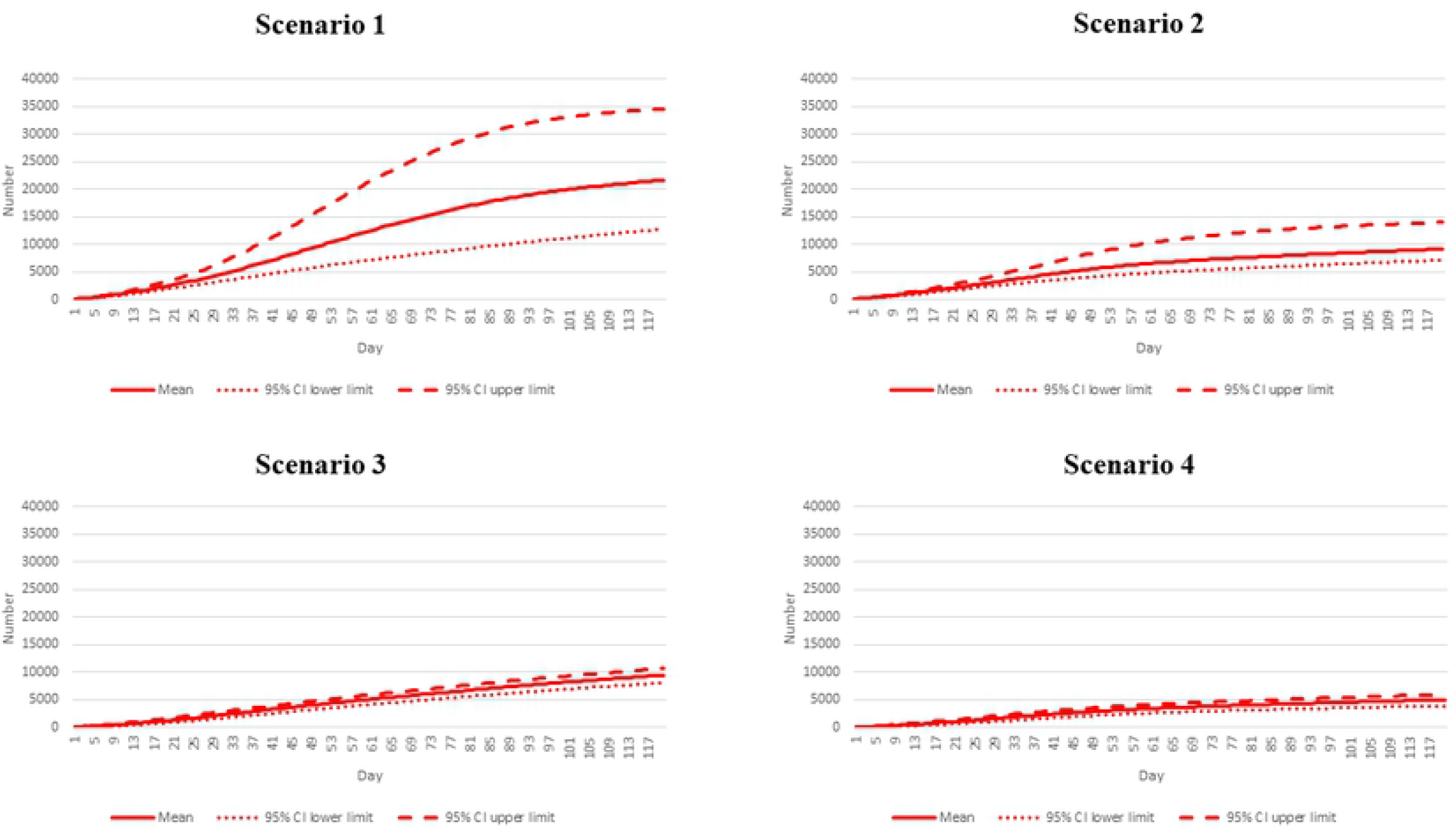
Cumulative death tolls by different epidemic scenarios.

## Discussion

Overall, the results suggest that the advent of the Omicron variant in Thailand may lead to a sharp increase in SARS-CoV-2 transmission. In the pessimistic scenario where the Omicron variant is very highly transmissible and the vaccination rate continues at the current pace, the peak incident cases may far exceed the 2021 Delta wave peak (~50,000 *versus* 22,000 cases a day). However, daily deaths due to Omicron epidemic may not outstrip the deaths in the Delta wave (~270 *versus* 300 deaths during the peak).

Our findings are in line with the current Omicron epidemic in numerous countries. The US has just faced a record high of daily new cases. About one million cases hit the US within a day in the beginning week of 2022 as hospitalization records approached 123,000, almost on par with the record high [32]. However, deaths remained fairly stable at about 1,400 a day, well below the previous epidemic [32]. The UK also experienced a rapid surge of the people infected with the Omicron variant, bringing the UK total cases during the first week of 2022 to almost 1.3 million, about 30% higher than the week before [33]. The fourth epidemic wave in South Africa, caused by the Omicron variant, exhibited a peak of about 23,000 daily new cases, about one fifth greater than the earlier peak in June 2021 [34].

Concerning policy implication, this research informs that although with a pessimistic assumption, the death toll may not exceed the prior peak, a large number of cases should not be overlooked because the volume of severe cases could skyrocket in proportion to the case toll. In the worst scenario, the need for ventilators almost reaches 2,500 people per day, by day 50. Such a demand has already surpassed the nationwide ventilator reserve, which is set about 2,200. Note that the reserve is for all patients needing invasive ventilators, not only COVID-19 cases. During the peak of the Delta wave, the prevalent intubated cases soared up to about 1,200 a day. With this demand, the Thai healthcare system was already stretched as most hospitals were in dire lack of intensive care beds and ventilators [35]. Another consideration is that the consumption of invasive ventilators by COVID-19 cases partly means a compromise in the quality of care for other patients [36, 37].

This study also affirms the value of vaccinations to fight the Omicron variant. The peak of daily cases in scenario 2 in which we assumed very high transmissibility of the Omicron variant is approximately 40% lower than the peak in scenario 1. Note that the benefit of vaccination diminishes when comparing scenario 4 with scenario 3 as the force of epidemic was presumed to be less severe. This finding aligns with latest evidence from the Canada and the UK, which suggests that, despite the immune-escape characteristic of the Omicron variant, the third dose of COVID-19 vaccine provides some protection at least in the immediate term [38, 39]. The bottom line is now Thailand is witnessing vaccine administration at about 320,000 doses a day, still far lower than the maximum capacity of 1,000,000 doses a day, the figure set by the Government as a campaign to beat the Delta wave last year [40]. Hence, the Government should consider expediting the national vaccination rate and at the same time communicate with its citizens to maintain a high degree of individual protective behaviors and emphasize the importance of NPI as part of the collective effort of society.

This study is subject to some limitations. First, we did not account for the granular difference in the transmission rate of the Omicron variant across provinces. Moreover, we did not consider the impact of localized interventions. Second, we postulated a constant proportion of severe cases over the course of analysis. This may not be the case as the case fatality rate or the proportion of patients encountering severe conditions may increase during the period of high strain of healthcare facilities. Third, the analysis was surrounded by numerous uncertainties of the assumptions and the parameters. Knowledge of the Omicron variant, both in virology or public health, is yet to be confirmed. Last, like in many other modelling studies, we de-constructed the force of epidemic into various components, including contact rate, immunization rate, and vaccine effectiveness, to identify public health implications. However, we realized that it is extremely difficult to disentangle the biological characteristics of the virus from the social interventions or exactly ascribe the epidemic phenomenon to a particular determinant. An obvious example is the R, which is not a biological constant of a pathogen as it is also influenced by many other determinants, such as environmental conditions and behaviors of the infectees. Thus, interpretation of the results should be made with caution.

## Conclusion

Under the most pessimistic presumption, the Omicron epidemic in Thailand may cause a peak of daily incident cases at about 50,000 by day 73. The peak daily death toll may reach 270 by day 50, corresponding with the peak prevalent intubated cases at about 2,200. The acceleration of vaccine rollout will push down daily cases by up to 40% and the death toll by up to 55%. These figures should be used as input for the planning of healthcare resources, especially intensive-care beds and respiratory ventilators to meet the high demand of care, given the coming Omicron epidemic. A national campaign to expedite vaccination rollout alongside an emphasis on the importance of individuals keeping a high protective guard is recommended.

## Acknowledgements

We are extremely grateful for the support of the DDC staff in terms of advice and data provision. We also thank the International Health Policy Program (IHPP), the MOPH, for the offer of office space and computational software. The interpretation and conclusions contained in this study are those of the authors alone and do not always reflect the view of the agencies the authors are affiliated with.

## Author contributions

**Conceptualization:** Rapeepong Suphanchaimat, Pard Teekasap

**Data curation:** Rapeepong Suphanchaimat, Mathudara Phaiyarom, Nisachol Cetthakrikul

**Formal analysis:** Rapeepong Suphanchaimat, Mathudara Phaiyarom, Nisachol Cetthakrikul

**Investigation:** Rapeepong Suphanchaimat, Natthaprang Nittayasoot

**Methodology:** Rapeepong Suphanchaimat, Pard Teekasap

**Project administration:** Rapeepong Suphanchaimat, Natthaprang Nittayasoot

**Visualization:** Mathudara Phaiyarom

**Writing – original draft:** Rapeepong Suphanchaimat, Natthaprang Nittayasoot, Mathudara Phaiyarom, Nisachol Cetthakrikul

**Writing – review & editing:** Rapeepong Suphanchaimat, Pard Teekasap, Natthaprang Nittayasoot, Mathudara Phaiyarom, Nisachol Cetthakrikul

## Supporting information

**S1 Fig. Percentage decrease of contact rate in relation to daily incident cases under the Omicron variant epidemic (base case)**

(DOCX)

**S2 Fig. Percentage decrease of contact rate in relation to daily incident cases under the Omicron variant epidemic (twice the rate of the base case)**

(DOCX)

**S3 Fig. Percentage decrease of contact rate in relation to daily incident cases under the Omicron variant epidemic (half the rate of the base case)**

(DOCX)

